# Disaggregation as an interaction mechanism among intestinal bacteria

**DOI:** 10.1101/2022.07.22.501191

**Authors:** Deepika Sundarraman, T. Jarrod Smith, Jade V. Z. Kast, Karen Guillemin, Raghuveer Parthasarathy

## Abstract

The gut microbiome contains hundreds of interacting species that together influence host health and development. The mechanisms by which intestinal microbes can interact, however, remain poorly mapped and are often modeled as spatially unstructured competitions for chemical resources. Recent imaging studies examining the zebrafish gut have shown that patterns of aggregation are central to bacterial population dynamics. In this study, we focus on bacterial species of genera *Aeromonas* and *Enterobacter*. Two zebrafish gut derived isolates, *Aeromonas* ZOR0001 (AE) and *Enterobacter* ZOR0014 (EN), when mono-associated with the host, are highly aggregated and located primarily in the intestinal midgut. An *Aeromonas* isolate derived from the commensal strain, *Aeromonas-MB4* (AE-MB4), differs from the parental strain in that it is composed mostly of planktonic cells localized to the anterior gut. When challenged by AE-MB4, clusters of EN rapidly fragment into non-motile, slow-growing, dispersed individual cells with overall abundance two orders of magnitude lower than the mono-association value. In the presence of a certain set of additional gut bacterial species, these effects on EN are dampened. In particular, if AE-MB4 invades an already established multi-species community, EN persists in the form of large aggregates. These observations reveal an unanticipated competition mechanism based on manipulation of bacterial spatial organization, namely dissolution of aggregates, and provide evidence that multi-species communities may facilitate stable intestinal co-existence.

**Significance:** The diverse microbial communities inhabiting vertebrate intestines influence many aspects of their hosts’ health and development. The rules governing community assembly and even the types of interactions that are possible among gut microbes remain poorly understood, however, with the role of spatial aggregation in interspecies dynamics being especially unclear. To address this, we performed live imaging studies of the larval zebrafish gut, focusing on two spatially distinct bacterial species that exhibit strong competition, discovering that the latter can induce a striking, rapid disintegration of aggregates of the former. This competition is attenuated in the presence of a particular set of additional bacterial species. Our findings reveal an unanticipated interaction mechanism and highlight that diverse communities may stabilize the gut microbiome.

## Introduction

The vertebrate gut is home to a diverse set of interacting microbial species, with microbiome composition correlating with aspects of host metabolism, digestion, immune response, development, and more [2, 11, 14, 17, 21, 29–31, 41, 53, 58, 63, 75, 76]. Dysbiosis of the gut microbiota has been linked to several diseases such as obesity, inflammatory bowel disease, and rheumatoid arthritis [4, 8, 18, 36, 42, 46, 54, 60, 70, 77]. Understanding the forces that shape intestinal communities is therefore of considerable importance, and many studies have explored how factors such as host diet and genetics can affect microbiome assembly and function. In general, spatial organization is a common and important aspect of microbial ecosystems; surface attached biofilms, microbial mats, and marine snow provide well-known examples [12, 23, 39, 45, 69]. This likely holds in the gut as well, but spatial information is usually difficult to obtain. Conventional approaches based on, for example, metagenomic sequencing of fecal samples are blind to structure, and most spatially resolved methods typically involve coarse sampling or fixation methods that can drastically perturb the features present in live animals [68]. Regarding inter-microbial interactions in particular, we know little about the mechanisms at play, which may include segregation to enable coexistence, contact-mediated competition, or more complex spatially-varying signals.

Larval zebrafish provide a model vertebrate system in which to observe spatial relationships among gut microbial species. The animals’ optical transparency and amenability to gnotobiotic techniques [7, 13, 33, 48], together with a library of fluorescently tagged native gut bacterial isolates [73], allows controlled experiments involving live imaging of bacterial communities [28, 40, 55, 57, 72]. Mapping the spatial structure of a number of gut bacterial species established that each, in mono-association with its host, has a characteristic fraction of its population forming dense, three-dimensional aggregates and is preferentially localized in particular regions of the gut [56]. Given the well-established ecological principle of competitive exclusion, by which species competing for the same limited resource cannot stably coexist, we might expect that if introduced together, two species with an affinity for the same region will strongly compete, leading to greatly reduced abundance for at least one of the two relative to the mono-association value. Spatial segregation of competitors, whether static or dynamic, provides a route to co-existence [1, 9, 22]; thus we might expect that reducing spatial overlap of intestinal species should weaken competition. How these principles may or may not be manifested in a real vertebrate gut remains unknown. As we show below, an example with a pair of zebrafish-native bacterial species confounds simple expectations.

Inter-species interactions are also influenced by timing, and the outcomes of competition can vary depending on whether species arrive simultaneously or not to an environment. Various ecological studies have shown that early colonizers can have an advantage over later species and can often resist invasion by late competitors [37, 59, 64]. Regarding the gut microbiome, the pre-existing community is believed to contribute to resistance to pathogen invasion [6, 65], though a greater number of controlled studies in animal models are needed to uncover underlying mechanisms [6, 35].

To illuminate possible connections between spatial organization and inter-bacterial interactions in the context of a vertebrate intestine, we performed a set of live-imaging-based studies of a strongly interacting bacterial species pair, using light sheet fluorescence microscopy to visualize bacterial locations and dynamics in situ. Prior work based on measuring species abundances, without spatial information, characterized the interactions between various zebrafish-commensal gut bacterial species and found several examples of strong competitive pairwise interactions, one such pair being *Aeromonas* ZOR0001 and *Enterobacter* ZOR0014 [66], each of which form dense aggregates in mono-association [56, 57, 72]. These and other strong pairwise interactions were dampened in the presence of additional bacterial species [66].

As described here, imaging reveals that both of these species colocalize and co-aggregate in the larval midgut. We introduce a derivative of *Aeromonas* ZOR0001, *Aeromonas-MB4* that is deficient in biofilm formation in vitro in the presence of a glycan commonly found in the gut. We find that in vivo, *Aeromonas-MB4* is largely planktonic and motile. The dispersed strain leads to an even lower *Enterobacter* abundance than does the co-localized wild type, and it induces a striking dissociation of *Enterobacter* aggregates. In the presence of an already established community of multiple bacterial species, competition and disaggregation effects are diminished, coexistence is enhanced, and the planktonic isolate behaves more similarly to the wild-type parental strain. Overall, our observations point to a surprising and unanticipated mechanism by which spatial structure can mediate gut bacterial interactions, namely induced disintegration of aggregates.

## Results

### Aeromonas-MB4 exhibits planktonic behavior in vitro and in vivo

We consider two strains of *Aeromonas*. One is a zebrafish gut commensal, *Aeromonas* ZOR0001 (referred to as AE). The other is an *Aeromonas* isolate (referred to as AE-MB4), derived through directed evolution of AE in vitro. The zebrafish-native AE forms aggregates in vitro in certain growth media, including sterile embryo medium supplanted with 0.4% N-Acetylglucosamine (GlcNAc), a sugar prevalent in intestinal mucus [43, 49, 51, 61]. The AE-MB4 strain was derived from AE by repeated passaging of fractions of such media that were devoid of visible aggregates, thereby selecting for mutant bacteria deficient in aggregation in the presence of GlcNAc (Fig S1). The molecular genetic characterization of AE-MB4 is described elsewhere [62]; we focus here on the spatial dynamics of this strain and the consequences of this altered aggregation behavior on community structure.

We first characterized the spatial distribution of bacteria in initially germ-free zebrafish larvae (Fig 1A) mono-associated with either the commensal AE or the AE-MB4 isolate. We mono-associated initially germ-free zebrafish at 5 days post-fertilization (dpf) with each strain and imaged 48 hours later using light sheet fluorescence microscopy (Methods). Wild type AE forms dense clusters in the midgut (Fig 1B, D), as seen in prior work [28, 55–57, 72] whereas AE-MB4 resides primarily in the anterior with a large fraction of the population in the form of discrete planktonic cells (Fig 1C,D, Video S1). Image analysis (see Methods) identifies individual bacterial cells and aggregated clusters and provides estimates of cluster populations. Cluster size statistics highlight the differences between the strains. Pooling cluster sizes from all fish, we compute the probability of a cell being in a cluster of a given size, i.e. the ratio of the number of cells found in a given cluster size bin to the total population from all clusters. For example, the bar associated with bin (10^0^ − 10^1^] reflects the frequency of finding cells in clusters of size 2-10 in the total population. We plot this measured frequency for different cluster size bins in Fig 1E. AE cells are most likely to reside in large clusters (n ≈ 10^3.5^), a population completely absent in AE-MB4. AE-MB4 cells are over 30 times more likely to be planktonic, i.e. in clusters of size 1, than AE. All cluster sizes are provided as supplemental Material (S2 File). We also plot the reverse cumulative probability distribution of cluster sizes, p(n), i.e. the probability that a given cluster is composed of more than n bacterial cells (Fig 1F). The distributions for both species have characteristic power law form for small clusters (n < 100) with a slope of m = 0.8 ± 0.1 [55]. The disparity in p(n) between AE and AE-MB4 grows as n increases and is especially pronounced for clusters of size n > 1000 as the distribution of AE becomes shallower. Earlier work studying intestinal cluster size distributions in a range of species has shown that characteristic power-law features arise from fragmentation and growth dynamics, and a shallow decline of p(n) at large n is indicative of aggregation [55], consistent with observations here.

**Figure 1.**
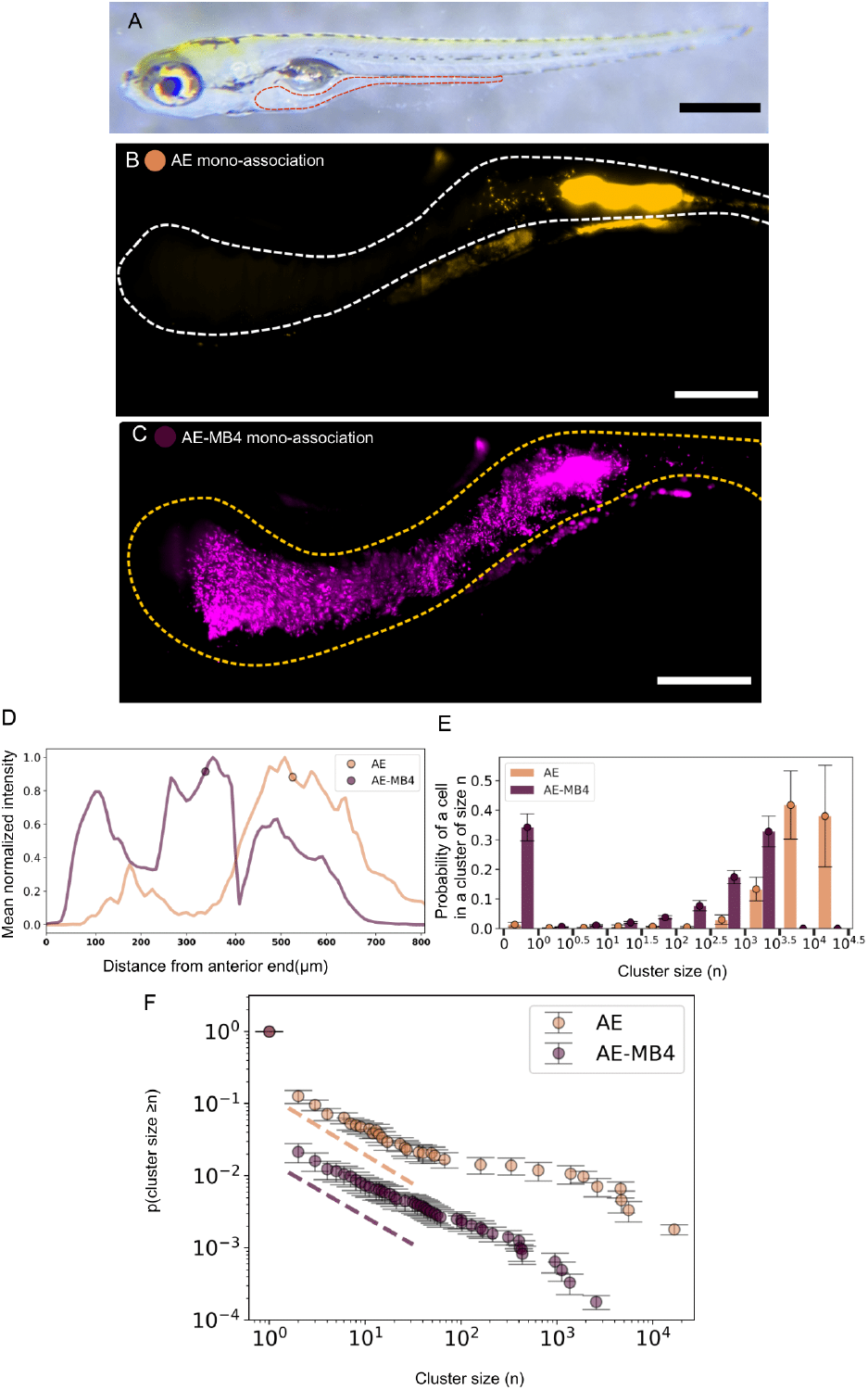
(A) Bright field image of a 7dpf zebrafish with the gut outlined in red. Bar: 500*µ*m (B & C) Maximum intensity projections of 3D images of the intestine of zebrafish mono-associated showing (B) zebrafish-commensal AE bacteria (orange), which forms large midgut aggregates and (C) AE-MB4 (magenta), which is largely planktonic. Dotted curves roughly indicate the gut boundary. Bar: 100*µ*m. (D) Mean normalized intensity profiles of AE (orange) and AE-MB4 (magenta) along the anterior-posterior axis. In contrast to AE, which localizes predominantly in the midgut, much of the AE-MB4 population is in the anterior bulb. Circles indicate the center of mass. Curves are averaged from N = 6 fish each. (E) The probability of a cell being in an n-cell cluster for AE (orange) and AE-MB4 (magenta), mono-associated with zebrafish. Circles and error bars indicate the mean and uncertainties, calculated using jack-knife resampling. Tick marks indicate bin intervals, e.g. the orange and magenta bars between 0 and 10^0^ correspond to AE and AE-MB4 probabilities to be in clusters of size n = (0-1]. Data are from N = 6 and 8 fish for AE and AE-MB4, respectively. (F) The cumulative cluster size distributions (p[cluster size ≥ n]) for AE (orange) and AE-MB4 (magenta) in mono-association calculated from all clusters found in N = 6 and 8 fish, respectively. showing power law behavior for n < 10^2^. The dashed lines illustrate the slopes from power law fits in the range n = 1 to 10^2^, with exponents 0.8 for each strain. At large n, the AE distribution plateaus, reflecting the presence of large clusters.

### A highly aggregated commensal species undergoes rapid fragmentation in the presence of the Aeromonas-MB4 isolate

To study the role of spatial coincidence in competition in the zebrafish gut, we studied interactions of *Aeromonas* and *Aeromonas-MB4* with a different zebrafish-commensal bacterial species, *Enterobacter* ZOR0014 (referred to as EN). Past work based on abundance measurements established that the parental AE strain has strong negative interactions with EN, with the population of the latter reduced by over an order of magnitude from its mono-association value when di-associated with AE (Fig 2E) [66]. EN in mono-association is almost completely aggregated, with only about 10% of its population in the form of planktonic individuals, forming large clusters primarily in the midgut region of the intestine (Fig 2A, Video S1) [56, 57]. Consistent with earlier measurements, its cluster size cumulative probability distribution and the probability of a cell being in a cluster of a given size reflect the highly aggregated spatial character of the species (Fig S2, S3).

**Figure 2.**
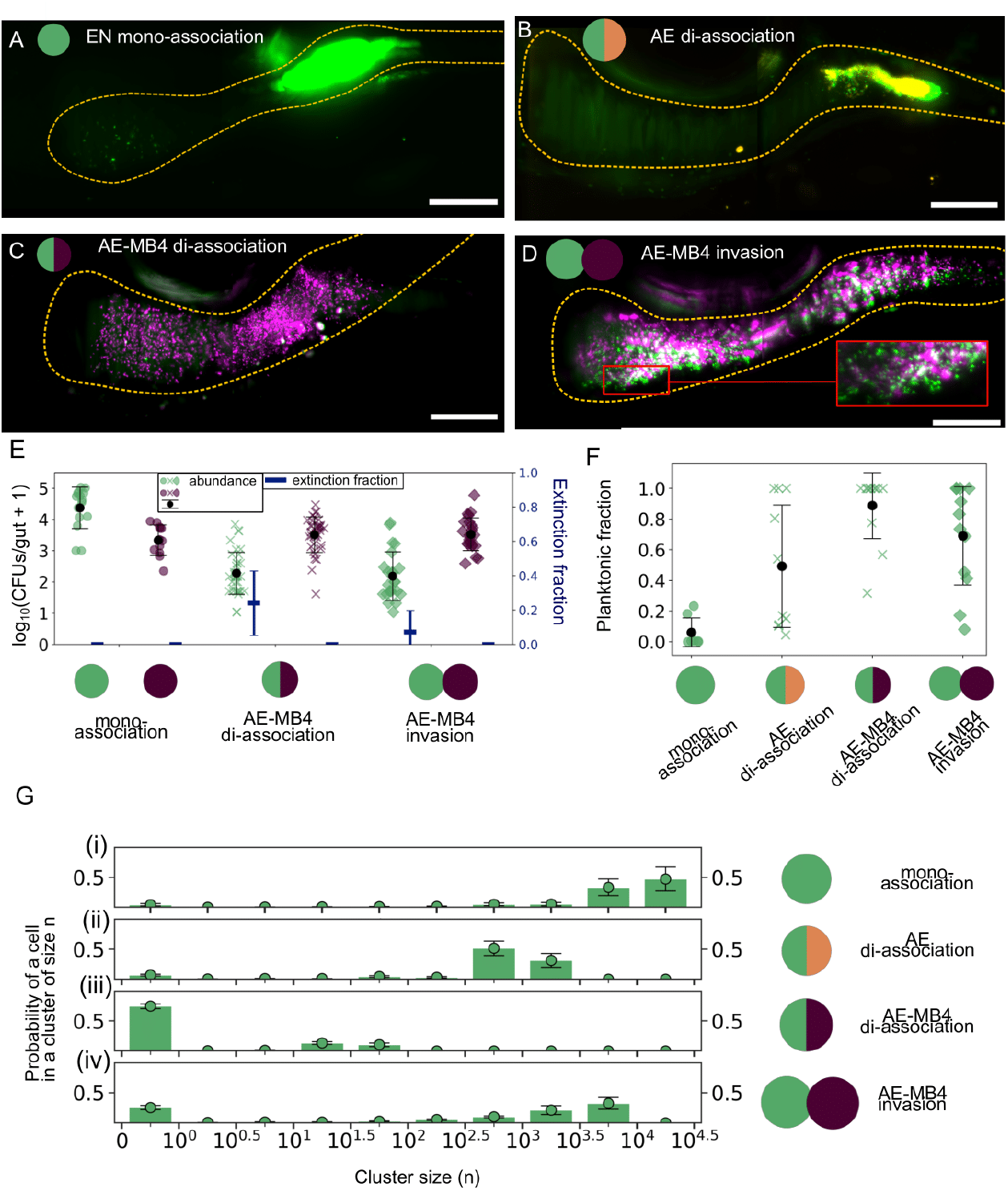
(A-D) Maximum intensity projections of 3D images showing (A) EN (green) in mono-association forming large clusters in the midgut, (B) EN (green) in di-association with AE (orange) showing co-aggregation of both species in the midgut, (C) EN (green) in di-association with AE-MB4 (magenta) showing sparse EN amid abundant planktonic AE-MB4 cells, and (D) EN (green) after invasion by AE-MB4 (magenta) showing large populations of single cells and small clusters of EN throughout the gut. Inset: Many single cells of EN are evident. Dotted curves roughly indicate the gut boundaries. (E) Twin axis plot showing the abundance (left axis) in log_10_(CFUs/gut + 1) and extinction fraction (right axis) for EN (green), AE (orange) and AE-MB4 (magenta) in mono-association (circles), di-association (Xs), and when EN is invaded by AE-MB4 (diamonds). Each marker indicates abundance data from a single fish. The blue line markers and black circular markers depict mean and standard deviation of the extinction fraction and abundance for EN and AE-MB4 in each of the experiments respectively. From left to right, N = 15, 12, 17, 27, 27, 36, 36, 23, and 23 fish. (F) The planktonic fraction per fish for EN in (from left to right) mono-association, dissociation with AE, dissociation with AE-MB4 and when invaded by AE-MB4. From left to right, N = 7, 10, 12, 12 fish. G. As in Fig. 1 E, The probability of a cell being in an n-cell cluster for EN (i) in mono-association, (ii) in di-association with AE, (iii) in di-association with AE-MB4, and (iv) invaded by AE-MB4. Circles and error bars indicate the mean and uncertainties. Data are from N = 7, 10, 12, and 12 for (i)-(iv), respectively.

We performed di-association experiments (as in [66]) in which both AE and EN were inoculated simultaneously at 5 dpf and the spatial distribution was assessed by imaging at 7 dpf. We found that AE and EN often co-aggregate in the same region of the intestine (Fig 2B, Video S2), suggesting that strong competition could be a consequence of both species having an affinity for similar locations. These co-aggregates of AE and EN consist of intermingled clonal clusters of each species, not well-mixed individuals of each species.

We next investigated the consequences of the AE-MB4 traits on the *Aeromonas-Enterobacter* competition. In di-association with AE-MB4, EN exhibits almost an order of magnitude lower mean abundance (2.2 ± 0.7) than with AE (3.0 ± 0.6), and roughly two orders of magnitude lower than its mono-association value (4.4 ± 0.7) (Fig 2E), in contrast to expectations that the species’ dissimilar localizations may attenuate competition. Extinction of EN was more frequent in di-association with AE-MB4 (extinction fraction = 0.2 ± 0.2), than with AE (extinction fraction = 0.1 ± 0.1). The effect on the spatial distribution of EN is striking; nearly all EN are found as discrete individuals (Fig 2C, Video S3), with the planktonic fraction being 0.9 ± 0.2 (mean ± std. dev.) compared to 0.1 ± 0.1 in mono-association. EN cells are never found in clusters of size n > 10^2^, the most common range in mono-association, and in nearly 80% of the fish examined, EN shows no apparent clustering at all. Individual cells of EN rarely co-aggregate with AE-MB4 clusters, in contrast to frequently occurring co-aggregates of both species in the di-association of EN with AE. All abundance data are provided in Supplemental File S1 and cluster sizes in Supplemental File S2.

To test how priority effects could impact interactions, we examined the abundance and spatial distribution of EN after invasion by either the AE or AE-MB4 strains. Prior work established that EN reaches steady state abundance in approximately 8-12 hours [57, 72]. We therefore inoculated the second species 24 hours after the introduction of EN. When challenged with AE, the initial advantage serves to benefit EN, with most of the fish sustaining EN aggregates (Fig S4, Supplemental File S1). We anticipated priority effects to similarly impact invasion by AE-MB4. Contrary to this expectation, we found that EN is unable to persist as large aggregates in the gut (Fig 2D, Video S4). Instead, the population consists of a large fraction of individual cells and some small clusters with a mean planktonic fraction of 0.7 ± 0.3 (Fig 2F,G), similar to that observed in the di-association experiments. The remaining clusters of EN in these experiments are often fragmented and sparse, unlike the dense aggregates of EN seen in other experiments.

Through time series imaging of EN-colonized fish starting 6 hrs after invasion with AE-MB4, we observed striking, rapid fragmentation of large EN aggregates into individual cells and small clusters within timescales of hours (Fig 3A,B, Video S4, S5, S6). Such disintegration has not been observed in mono-association or di-association with AE and thus we attribute it to the presence of AE-MB4. The planktonic EN showed little motility (Video S7). We identified in images planktonic and aggregated sub-populations of EN following invasion and separately calculated the growth rate (r) of each from exponential fits of abundance as a function of time (Methods), considering only regions in which there was growth, not influx or expulsion of bacteria. We found the growth rate of individual cells (r = 0.24 ± 0.11 hr^−1^, mean ± std. dev. for N=3 fish) to be approximately three times lower than that of clusters (r = 0.66 ± 0.23 hr^−1^ for N=4 fish) (Fig 3C). The large planktonic fraction together with the lower growth rate of the planktonic relative to the aggregated form implies a reduced ability of EN to persist in the gut following AE-MB4 invasion. We also assessed from images the growth rate of AE-MB4 in the invasion experiments. Its value (r = 0.76 ± 0.04 hr^−1^ for N=3 fish) was higher than that of EN, consistent with the expectation that a fast growing species outcompetes a slow growing species.

**Figure 3.**
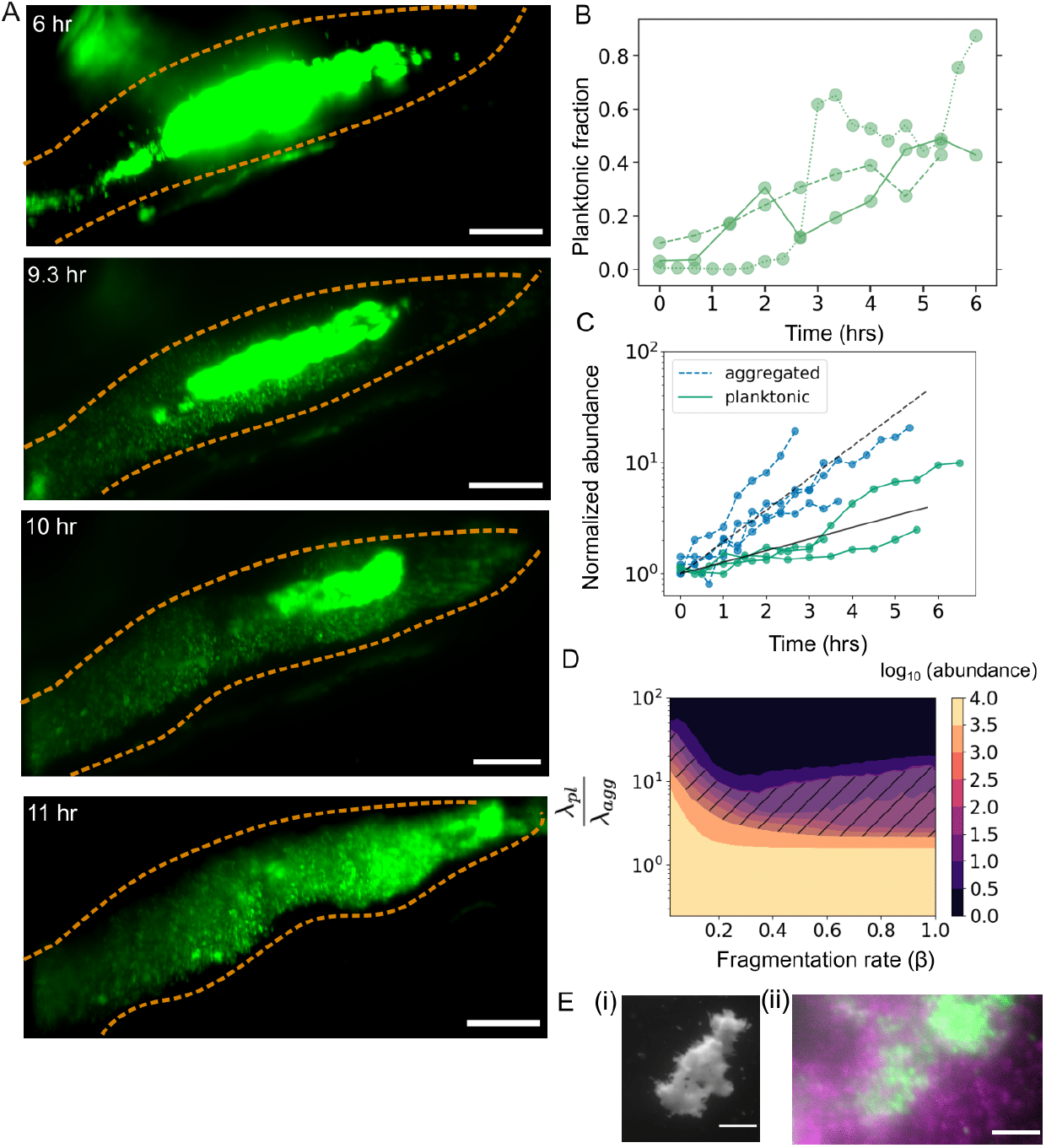
(A) Maximum intensity projections of the midgut region from a time series showing an aggregate of EN that fragments following invasion by AE-MB4. Times indicate hours post-invasion. Dotted curves roughly indicate the gut boundary. Bar : 50 *µ*m. (B) The planktonic fraction of the EN population over time for 3 fish measured starting 6 hrs. after invasion by AE-MB4. Data provided in Supplemental File S6. (C) The number of cells normalized by the initial value over time for aggregated (blue, dotted) and planktonic (teal, solid) EN populations after invasion by AE-MB4. For aggregated populations, each dataset depicts the growth of a single cluster of EN in a distinct fish (N=4). For planktonic populations, each dataset corresponds to a region comprising only planktonic cells in a distinct fish (N=3). The black curves indicate the mean growth rates: m = 0.66 ± 0.23 hr^−1^ and 0.24 ± 0.11 hr^−1^ for aggregated and planktonic cells, respectively. (D) Simulated abundance of EN as a function of the ratio of the expulsion rates for planktonic to aggregated populations 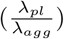 and the fragmentation rate *β* (hr^−1^). Experimentally measured parameters were used for the growth rates of aggregated (*r*_*agg*_) and planktonic (*r*_*pl*_) EN clusters (shown in C). The shaded region corresponds to the experimentally measured abundance (log_10_(abundance) = 2.0 ± 0.9). See the main text for details. (E) Brightfield image of EN-AE-MB4 co-aggregates in 0.4 % GlcNAc solution. Scale bar: 1mm. (ii) Single plane of a 3D image showing co-aggregated populations of AE-MB4 (magenta), EN (green) and overlapping populations (white) in 0.4 % GlcNAc. Scale bar: 10*µ*m.

To better understand the implications of a high planktonic fraction of slow-growing, non-motile EN in the gut, we simulated the dynamics of clusters in the zebrafish intestine undergoing four key processes, growth, aggregation, fragmentation and expulsion. Simulation of these processes has been shown to reproduce characteristic features of cluster size distributions observed in experimental data [55].

A detailed description of the stochastic model and its parameters is provided in Methods. In brief, growth of each cluster is logistic and the total abundance (the sum of all cluster populations) is bounded by a carrying capacity, K, equal to the mono-association abundance of the species. Growth rates of individuals and clusters (r_pl_ and r_agg_) are set to the experimentally measured values (Fig 3C). Aggregation, i.e.the combining of two clusters of size n and m to form a single cluster of size n + m, occurs at a rate *α* _nm_ = *α* (n m)^1/3^, where the exponent 1/3 signifies a collision likelihood proportional to cluster diameter. The value of the rate prefactor *α* is based on prior work: log_10_(*α*) = −2.5. A cluster of size n undergoes size-dependent fragmentation of single cells at a rate *β* = (*n*)^2/3^, with the 2/3 exponent reflecting that only cells on the surface leave a cluster. The value of *β* is varied in simulations, over a range corresponding to 2-100 cells fragmenting per hour from a cluster of size 1000.Size-independent expulsion of aggregated clusters is fixed at a rate set by previous studies, λ_agg_=0.1 hr^−1^, while the expulsion rate of planktonic cells is varied between λ _pl_ = 0.025 and 10 hr^−1^.

We combine these processes in a stochastic simulation. At each time step, one of the fragmentation, aggregation, or expulsion reactions occurs with a probability determined by the rates for each reaction, computed for each cluster. First, the simulation is run to determine the mono-association cluster distribution of EN for fixed *α, β* and λ after 24 hrs. After generating this initial EN distribution, we simulate the AE-MB4 invasion of EN aggregates by varying the fragmentation rate, *β*. We run this for a range of values of λ_pl_. Averaging the total abundance from 500 replicates for each pair of expulsion rate ratio 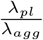 and fragmentation rate *β* parameters we obtain the abundance phase diagram shown in (Fig 3D). When λ_*pl*_ *< λ*_*agg*_, we find that abundance is similar to the mono-association value, irrespective of fragmentation rate. For λ_*pl*_ > λ_*agg*_, the abundance corresponds to the experimentally observed post-invasion abundance values for an expulsion rate 0.13 hr^−1^ < λ_*pl*_ < 2.5 hr^−1^ with the overlap more distinct in the high fragmentation regime. This is physically plausible as non-motile cells are passively driven by intestinal flows and hence should have a large expulsion rate. This suggests that the properties of planktonic, non-motile EN, with its reduced growth rate and increased susceptibility to being expelled from the gut, can explain the low experimentally observed values of EN abundance.

We also studied the aggregation of EN in the presence of AE-MB4 in in vitro experiments, examining the bacteria in media with 0.4 % GlcNAc, in which AE-MB4 resists aggregation (Fig S1). Strikingly, in co-culture experiments of AE-MB4 and EN in this medium, AE-MB4 readily co-aggregates with EN and we observed large clusters comprising both species Fig. 3E. These observations show that in vivo interaction dynamics and outcomes are markedly different from those in vitro.

### The Aeromonas-MB4 isolate displaces the closely related parental AE strain

To test whether the strong competition and dynamics observed are specific to EN and to examine how other gut bacterial species may be impacted by AE-MB4, we studied AE-MB4’s interactions in the gut with the parental zebrafish isolate AE strain. We examined intra-species interactions by inoculating, as in previous experiments, initially germ-free zebrafish with fluorescently labeled AE at 5 dpf and challenging by introduction of either AE-MB4 or differently-labeled AE 24 hours later. The abundance of AE after invasion by AE-MB4 (log_10_(abundance) = 2.8 ± 0.5) drops by over an order of magnitude compared to its mono-association value (log_10_(abundance) = 4.2 ± 0.4) (File S1). Comparing the probabilities of finding AE cells in different sized clusters when invaded by itself versus when invaded with the planktonic AE-MB4, we discovered an absence of very large clusters (n = 10^4^) in the presence of AE-MB4 (Fig 4, Video S8,S9), suggesting that the AE-MB4 isolate impacts the parental strain’s ability to either form or maintain dense aggregates. The dynamics of interactions differ, in that we observe no dissociation of AE aggregates and instead expulsion of large AE clusters with often only small clumps persisting (Video S10).

**Figure 4.**
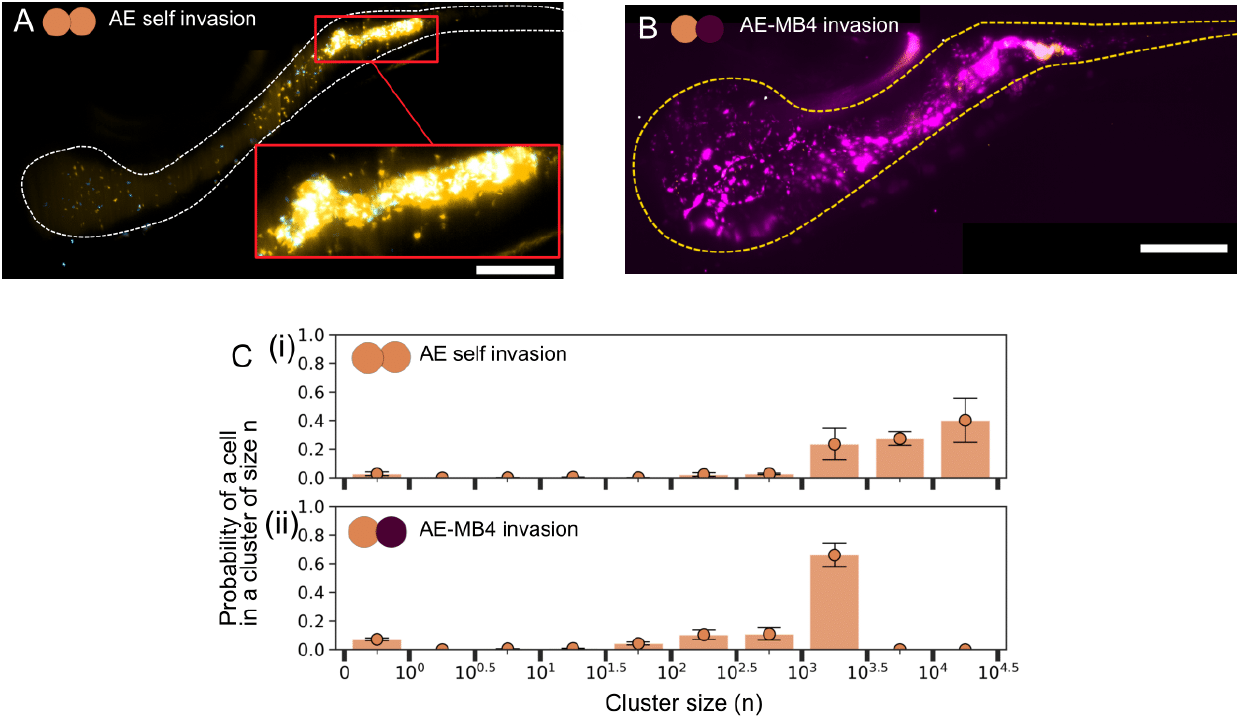
(A & B) Maximum intensity projections from zebrafish initially colonized by AE (orange), invaded 24 hrs. later by (A) differently labeled but otherwise isogenic AE (blue) (B) AE-MB4 (magenta). Images (A) and (B) were taken approximately 24 hrs. post-invasion. Dotted curves roughly indicate the gut boundary. Bar: 100*µ*m. Inset to (A): a bacterial aggregate comprising both populations of AE (white overlapping regions). (C) Cluster probabilities as in As in Fig. 1E for AE (orange) when (i) invaded by others of the same strain (AE) and (ii) invaded by AE-MB4, both after 24 hrs. Circles and error bars indicate the mean and uncertainties. Data are from N = 8 and 10 fish for (i) and (ii), respectively.

### Multi-species communities dampen strong AE-MB4-EN interactions

We next studied the interactions of EN and AE-MB4 in the presence of additional species, using a set of zebrafish commensal bacteria whose interactions have previously been characterized [66]. The additional gut community members comprised *Plesiomonas* ZOR0011 (referred to as PL), *Acinetobacter* ZOR0008 (AC), *Pseudomonas* (ZWU0006) (PS), and *Aeromonas* (AE). First, we co-inoculated four species (EN, AC, PL and PS) at 5 dpf and observed high abundances and large populations of aggregated EN in the midgut region of the intestine (Fig 5A), resembling its mono-association form (Fig 2A, E, G, Video S11), suggesting that these species have little effect on Enterobacter’s spatial structure. Then we examined the impact of AE-MB4 when also co-inoculated along with the other four species and found that on average populations of EN were more planktonic (planktonic fraction = 0.7 ± 0.2), (Fig 5B, Video S12), as seen in the di-association experiments. We also noted frequent extinction of EN, with approximately 1/3 of the fish showing no trace of EN. Overall, however, EN abundance was slightly higher (log_10_(abundance) = 2.8 ± 0.5) than in the di-association experiments (log_10_(abundance) = 2.3 ± 0.7) (Fig 5E,G). This suggested that the negative AE-MB4 interaction effects are still present but relatively weaker in these five species co-inoculation experiments. The abundance data (Fig 5E) includes fish in which all species inoculated in the experiment were detected and all species excluding PL were detected. PL generally showed rare colonization across all datasets.

**Figure 5.**
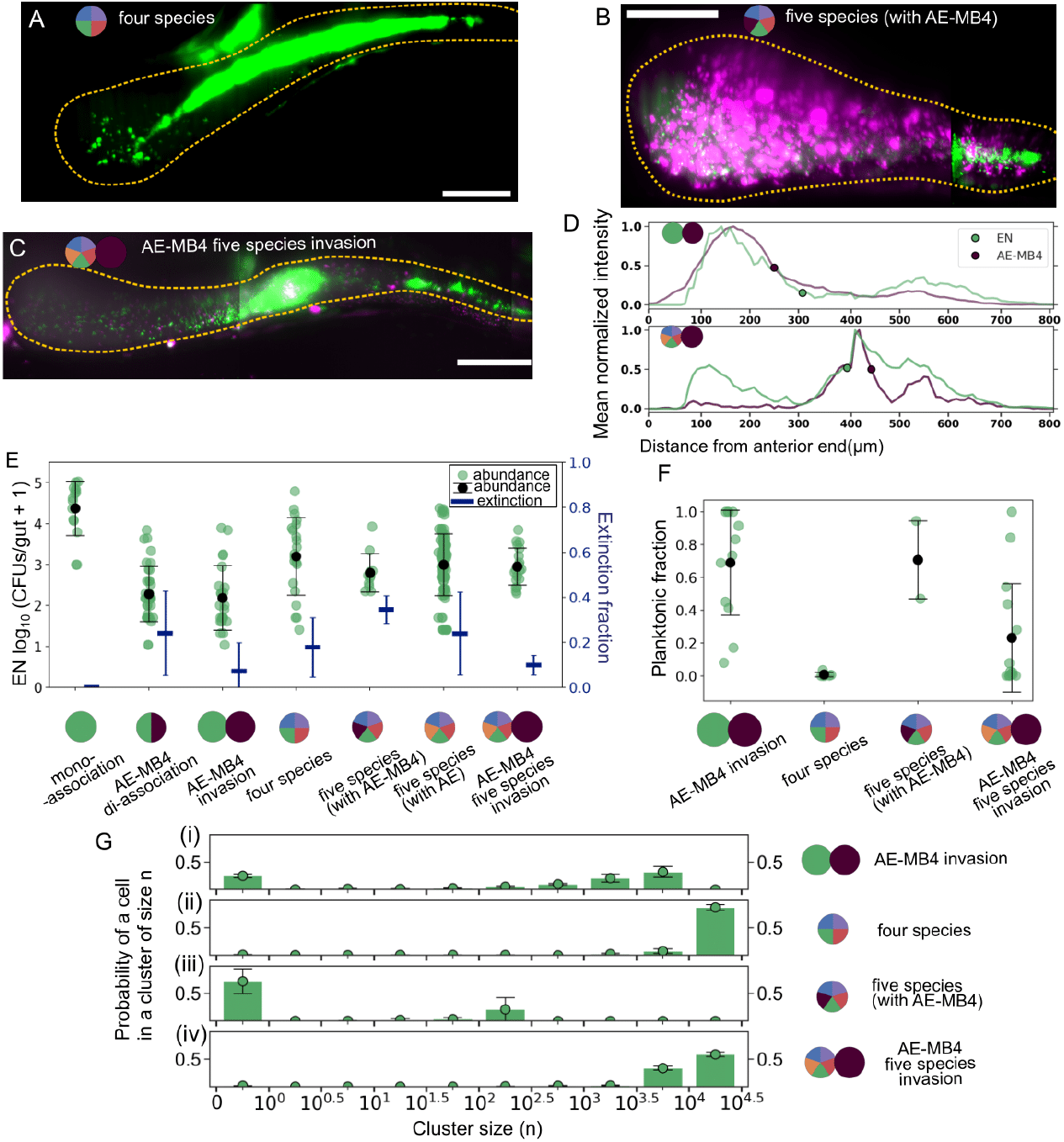
Maximum intensity projections of a larval zebrafish gut showing the spatial distribution of (A) EN (green) in the presence of four other commensal gut bacterial species forming large aggregates in the intestine (B) EN (green) in the five-species co-inoculation experiment (including AE-MB4 (magenta)) has smaller fragmented clusters and many single cells (C) EN (green) in the experiment when five species (including AE) are invaded by AE-MB4 (magenta) where EN forms larger clusters with some single cells and AE-MB4 populations show some aggregation. The dotted curve roughly defines the gut wall. Bar:100*µ*m (D) Mean normalized intensity profiles measured along the anterior-posterior axis showing the spatial distribution of EN (green) and AE-MB4 (magenta) along the intestine when (i) EN is invaded by AE-MB4 and (ii) all five species are invaded by AE-MB4 showing large aggregates of EN persist in the midgut. The circular markers indicate the center of mass. Data are from N = 13 and 11 fish for (i) and (ii) respectively. (E) Twin axis plot showing the abundance of EN (left axis) in log_10_(CFUs/gut + 1) and extinction fraction (right axis) when in mono-association, in di-association with AE-MB4, invaded by AE-MB4, in the four-species co-inoculation experiment, in the five species co-inoculation experiment (including AE-MB4), in the five species co-inoculation experiment (including AE) invaded by AE-MB4. Abundances from individual fish are depicted with green circular markers. Mean and standard deviation for abundance and extinction fraction of EN are shown with black circular and blue line markers respectively. N (from left to right) = 15, 36, 23, 21,11, 87, 19 fish. (F) The planktonic fraction per fish for EN when invaded by AE-MB4, in the four–species co-inoculation experiment, in the five-species co-inoculation experiment (including AE-MB4) and when the five-species (including AE) are invaded by AE-MB4. Black markers indicate the mean and standard deviation. N (from left to right) = [12, 5, 2, 14] fish. (G) The probability of being in an n-cell cluster of EN when (i) invaded by AE-MB4 (ii) in the four-species co-inoculation experiment (iii) in the five-species co-inoculation experiment (including AE-MB4) (iv) in the five-species co-inoculation experiment (including AE) invaded by AE-MB4. Mean and uncertainties determined from jack-knife resampling are shown with black markers. N (top to bottom) = [12, 5, 2,14] fish.

To determine whether these interactions persist in spite of priority effects, we performed invasion experiments with fish co-inoculated with all five species (AC, AE, EN, PL and PS), similar to mono-association EN experiments. We first studied the abundance distribution of EN in the control experiment, when no invader was introduced. We found that EN is more likely to be aggregated in these fish, resembling its typical structure with mean abundance 10^3.0±0.8^ (Fig 5E, Video S14).

On challenging this multi-species community with AE-MB4, we found strikingly different results from that of the mono-association invasion. Large aggregates of EN were able to persist in the presence of other species and the mean planktonic fraction of EN (0.2 ± 0.3) was lower than in the mono-association invasion experiments (0.7 ± 0.3) (Fig 5 D,C and F). The likelihood of a cell being in clusters of size n > 10^3^ goes up from 67% in the mono-association invasion experiment to 97% in the multi-species challenge experiments (Fig 5G), while the planktonic probability is reduced from 25% to 2%. In time series experiments observing the dynamics of EN aggregates in the presence of other species, large aggregates of EN either resist breakup or undergo slower fragmentation relative to the fast timescales observed in single species mono-association invasion experiments (Video S16). Together, all of these observations suggest that a diverse and stable pre-existing community can inhibit strong interactions with the invader.

To quantify the strength of interactions in different contexts, we calculated an interaction coefficient measuring the effect of species j on species i and defined as follows as in [66]:

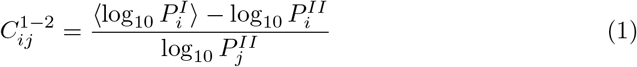

Here, 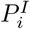 is the mono-association abundance of species i, and 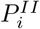 and 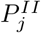 are the abundances of species i and j in di-association. This follows from a competitive Lotka-Volterra model using log-transformed abundances in place of absolute abundance, described in detail in [66].

One can extend this interaction coefficient to quantify the effect of a single species j on another species i in the five species experiments:

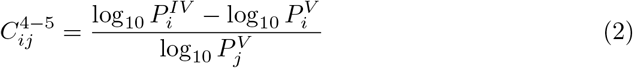

These equations can be applied to the invasion experiments where 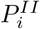 and 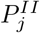 are replaced by the abundances of species i and j in the invasion experiments. Similarly, Eq.2 translated to the five species invasion case can be written as:

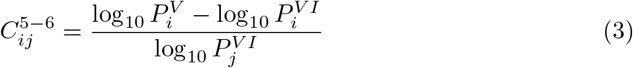

where 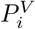 is the abundance of species i in the five species (AC, AE, EN, PL and PS) co-inoculation experiment, 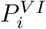 is the population of i post-invasion with species j and 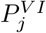 is the abundance of the invader j. Using equations Eq. 1, 2 and 3, with the abundance data provided in File S1, we can extract the mean and uncertainty of the interaction coefficients for various experiments. We measured the interaction coefficients in the di-association and invasion experiments to be similar with 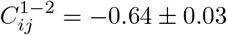 for the di-association and −0.65 ± 0.03 for the invasion experiments. These signify strong negative interactions, with EN abundance dropping by two orders of magnitude relative to mono-association. In the 5 species co-inoculation experiments with AE-MB4, we found 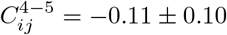, significantly weaker than 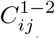 for di-association. For the invasion experiments we found the magnitude of interactions to be zero within uncertainties 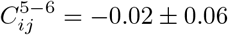. These coefficients affirm implications of previous work that higher order interactions dampen strong pairwise effects and promote co-existence in a diverse community.

To further examine the nature of interactions in a multi-species context, we returned to characterizing the effects of AE-MB4 on the parental AE strain. In mono-association, challenged by AE-MB4, AE was unable to persist in clusters of n ≥ 10^4^ cells with no likelihood of being in such clusters (Fig 4C (ii)). When additional species are present, we see large aggregated populations of AE in the midgut and cells are 53% likely to be found in aggregates of size ≥ 10^4^ cells (Fig 6A, B, C, Video S15). As with EN, greater gut community diversity counteracts the changes induced by the AE-MB4 isolate. Surprisingly, we also found that AE-MB4, highly planktonic on its own, forms aggregates in the invasion experiment with other species present, also altering its localization in the intestine ((Fig 6B(ii)D).

**Figure 6.**
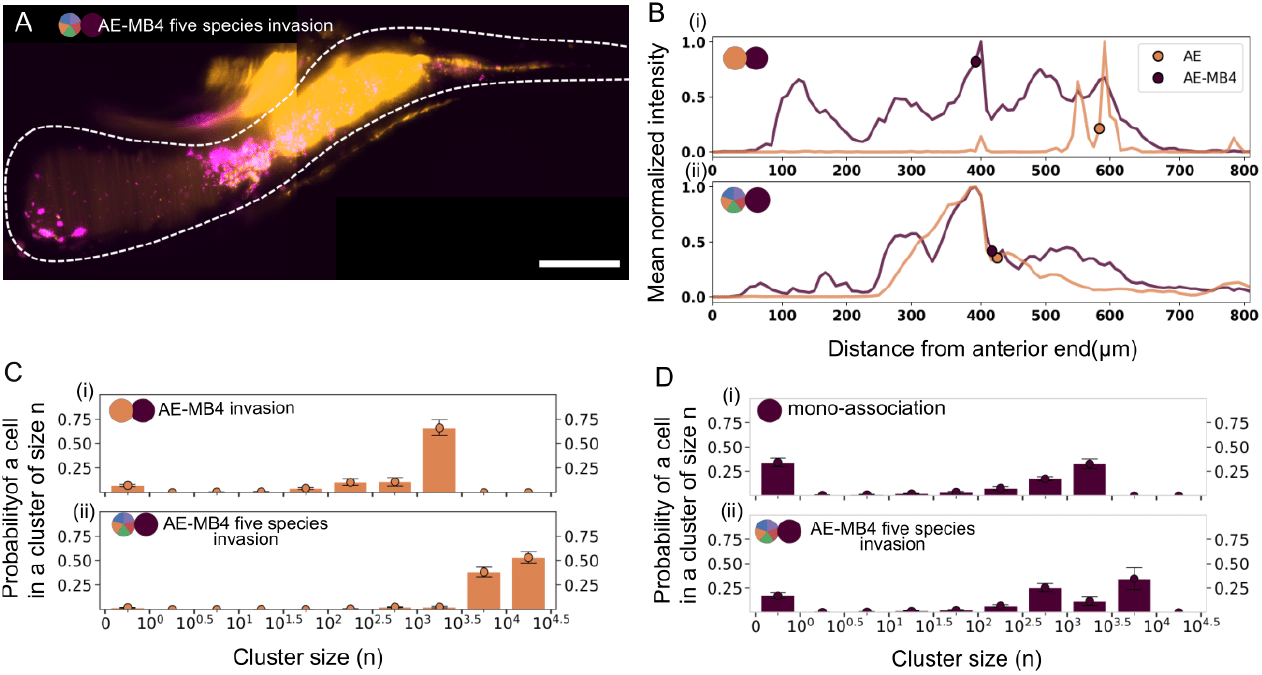
(A) Maximum intensity projection of a larval zebrafish gut showing a large cluster of AE (orange) after invasion by AE-MB4 (magenta) in the presence of the other four species. Small clumps of AE-MB4 are interspersed in the AE clusters. Bar: 100*µ*m (B) Mean normalized intensity profiles measured along the anterior-posterior axis showing the spatial distribution of AE (orange) and AE-MB4 (magenta) along the intestine when (i) AE and (ii) all five species (including AE) are invaded by AE-MB4. The circular marker indicates the center of mass. Data for (i) and (ii) are from N = 10 and 11 fish respectively. (C) The probability of being in an n-cell cluster for AE when (i) invaded by AE-MB4 (ii) invaded by AE-MB4 in the presence of the other four species. The mean and standard deviation are indicated with circular markers. The x tick marks indicate the bin intervals i.e. the bar between the tick marks 0 and 10^0^ corresponds to the probability of being in a cluster within (0-1]. Data are from N = 10 and 14 fish for (i) and (ii) respectively. (D) The probabilities (as in C) for AE-MB4 (i) in mono-association (ii) after invading the five species. The mean and standard deviation are indicated with circular markers. N = 8 and 11 fish for (i) and (ii) respectively.

## Discussion

We report an unexpected mechanism of interactions among gut bacterial species: induced disaggregation. Specifically, in our experiments, the presence of a non-aggregating strain disrupts a different, normally aggregated, commensal species, lowering its abundance and growth rate and altering its spatial structure in the intestine. Although the AE-MB4 isolate has reduced spatial overlap with EN than the parental strain, we find that this does not attenuate competition, defying the simple expectation that species with different localization characteristics are likely to have weaker interactions.

The molecular mechanism of the observed disaggregation is currently unknown. As described in detail elsewhere [62], the *Aeromonas-MB4* isolate investigated here was derived by directed evolution in media rich in N-Acetylglucosamine (GlcNAc), a sugar prevalent in intestinal mucus [15, 20, 25, 32]. *Aeromonas* is normally highly aggregated in the zebrafish gut, with clusters likely held together by extracellular polysaccharides and proteins as is typical for biofilms and other microbial aggregates. It is therefore plausible that in the AE-MB4 isolate, intestinal cues that inhibit production of polysaccharide-digesting enzymes are ignored, and these enzymes act broadly to degrade the extracellular matrix of *Enterobacter*. Testing this hypothesis requires considerable further study and could benefit from experimental assays such as using microfluidics to identify physical, chemical, and geometric conditions in which dissociation occurs. Similar phenomena, however, have been reported in non-intestinal contexts. For example, *Streptococcus pyogenes* secretes a protease that disrupts *Staphylococcus aureus* biofilms [5], the algae *C. reinhardtii* can inhibit aggregation of *Escherichia coli* [34], and mucus enhances in vitro biofilm formation by Escherichia coli [47] Given the observed co-aggregation of both wild-type *Aeromonas* and *Enterobacter*, it is likely that both species share cues and the disruption of aggregation of one, may directly interfere with aggregation of the other. Universal cues for inter-species bacterial communication such as autoinducer-2 have been known to promote formation of mixed species biofilms [10, 38]. A study has found that the suppression of autoinducer-2 expression in one of the species in a dual-species biofilm resulted in sparse co-aggregation and decreased biomass [50].

The *Aeromonas/Enterobacter* interactions we observe in the zebrafish gut differ strikingly from interactions observed in vitro. In minimal media with 0.4% GlcNaC, we find co-aggregates of AE-MB4 and EN (Fig 3E). yellowThe species intermingle at the scale of single cells, a morphology never seen in the gut. The dissimilarity is not surprising, as both the chemical and physical environment of the intestine differ from that of a media-filled petri dish, but it highlights the challenges of inferring in vivo behavior from in vitro assays.

The disaggregation and low abundance induced in *Enterobacter* by *Aeromonas-MB4* in pairwise competition in the zebrafish gut are attenuated in the presence of additional bacterial species. The ability of such higher order interactions to stabilize diverse communities has been noted in many contexts [3, 27, 44, 66], and the observations described here suggest that maintenance of spatial structure is a means by which multi-species coexistence is manifested. It is not currently clear whether this maintenance is facilitated by the biochemical activity of specific members of the community or by more complex spatial organization of species that may emerge. Studies involving labeled subsets of these species, or mutants derived with different aggregation traits, will illuminate the mechanism.

We also note the role of priority effects in determining community composition. It is known that early gut microbiome composition in humans influences whether a new colonizer can establish itself [65]. Studies have linked a diverse and healthy gut microbiome to colonization resistance [26], for example from invasion by the pathogen *Clostridioides difficile* [24, 67]. In the larval zebrafish, we find that in the presence of a diverse pre-existing community, AE-MB4 interactions with EN and AE are significantly weaker than when all species are inoculated simultaneously. Imaging reveals the spatial signatures that distinguish the outcomes of co-inoculation and invasion: AE-MB4 is more planktonic and less aggregated in the former than the latter, and EN maintains its aggregated phenotype with little fragmentation in invasion but not in co-inoculation studies.

The presence of gut bacterial aggregates [19, 55, 56, 72] and their importance to phenomena such as antibiotic response [57] and resistance to intestinal contractions [28] have been deduced in past work involving live imaging in larval zebrafish. This model system enables exploration of the community-level consequences of altered aggregation of a commensal bacterial species, lending insight into biophysical and biochemical mechanisms that may be at play more broadly, for example in the human gut. We suggest that bacterial manipulation of aggregation state, both their own state and the state of other species, may be a common feature of gut microbiome dynamics. Moreover, controlled disruption of bacterial aggregates could have therapeutic applications. An engineered disruptor, we speculate, could be introduced to the human gut to displace resident, aggregated microbes, facilitating their replacement by transplanted or otherwise intentionally provided strains. Reaching this end will require a more thorough understanding of inter-species modulation of physical structure.

## Materials and Methods

### Animal Care

All experiments with zebrafish were performed in accordance with protocols approved by the University of Oregon Institutional Animal Care and Use Committee and by following standard protocols [71]

### Gnotobiology

Wild-type (AB X TU line) larval zebrafish (*Danio rerio*) were derived using protocols described in [33]. The larvae at 0 dpf are washed with antibiotic, bleach and iodine solutions and then moved to tissue culture flasks containing sterile embryo medium solution at a density of approximately 1mL/larva. The flasks are stored in a temperature-controlled room maintained at 28C.

### Bacterial strains

Zebrafish gut isolates were previously tagged with green fluorescent protein (GFP) and dTomato [74]. The various isolates used for this study are textitAeromonas sp. (ZOR0001), *Enterobacter sp*. (ZOR0014), *Plesiomonas sp*. (ZOR0011) *Pseudomonas mendocina* (ZWU0006) and *Acinetobacter calcoaceticus* (ZOR0008). *Aeromonas-MB4* strain (JS168), an isolate of *Aeromonas* strain (ZOR0001) was generated through a guided evolution experiment described in [62]. All bacterial stocks were made in 25% glycerol and maintained at −80^*◦*^C.

### Inoculation of tissue culture flasks

The bacteria from frozen stocks were grown in lysogeny broth (LB medium) and shaken overnight for approximately 16 hrs in a temperature controlled room at 30^*◦*^C. 1mL of the overnight culture was washed twice by centrifuging at 7000g for 2 min and replacing the supernatant with fresh sterile embryo medium each time. For mono-association and co-inoculation experiments, the species were inoculated in fish at 5dpf and for the challenge experiments, the first species was inoculated at 5dpf and the second species is inoculated approximately 24 hrs after, at 6 dpf such that the flask water concentration of species is approximately 10^6^ CFUs/mL.

### Gut dissections and plating

We determined the abundances of bacterial species in the competition experiments by dissecting and plating the intestine. At 7dpf, zebrafish were euthanized via hypothermic shock, following which gut dissections were done to isolate intestinal contents. The intestine was placed in 500*µ*L of sterile embryo medium and homogenized by blending with 0.5mm zirconium oxide beads. Dilutions of 10^−1^ and 10^−2^ were prepared and 100*µ* L of these solutions were spread on tryptic soy agar plates. For experiments with zebrafish inoculated with more than 2 species, guts were plated on Universal HiChrome agar (Sigma Aldrich) to distinguish species based on a colorimetric indicator. The CFUs on the plates were counted to ascertain the abundances of the inoculated bacterial species. This data for each of the experiments is provided in File S1.

### Live Imaging using light sheet fluorescence microscopy

3D imaging of the larval zebrafish gut was done on a home-built setup described in [19]. The larvae were anesthetized with MS-222 (Syndel) and mounted in 0.6% agarose gel in glass capillaries. Each capillary is inserted into an imaging chamber containing sterile embryo medium and MS-222 at a concentration of 20*µ*L/mL of embryo medium. Once mounted, the fish were extruded from the capillaries. Lasers at 488nm and 568nm were used to excite GFP and dTomato bacteria respectively. Image acquisition involves translating the specimen in z-steps of 1 micron to capture the entire volume of the zebrafish gut. For time series experiments, imaging was done overnight at intervals of 20 or 30 minutes. Each acquired image comprises 2-4 regions that are stitched together using software. After imaging, the fish were euthanized by placing in an MS-222 solution of 40*µ*L/mL of sterile embryo medium.

### In vitro aggregation assay

AE and EN bacterial cultures were grown over night with shaking at 30^*◦*^C, normalized to OD600=5.0, then washed 2X in equal volume buffer (sterile embryo medium).

Normalized AE and EN cell suspensions were then back diluted 1:10 into 12-well plates containing buffer supplemented with 0.4% GlcNAc and incubated for approximately 6 hours with gentle rotation (115 RPM) at 30^*◦*^C to allow co-aggregate formation.

### Simulation

The model of bacterial cluster dynamics applied to EN is based on earlier work investigating bacterial cluster size distributions in the zebrafish gut [55]. We combine four kinetic processes – growth, aggregation, fragmentation, and expulsion – in a stochastic model simulated with a Gillespie algorithm. Growth is deterministic and occurs at every time step while aggregation, fragmentation and expulsion are stochastic. Each process is described by a kernel with specific rate parameters described below.

#### Growth

We model growth of each cluster as deterministic and following a simple logistic growth curve, with the total abundance of all clusters barred from exceeding an overall carrying capacity. The carrying capacity was fixed at 10^4^ cells, approximately the mono-association abundance of EN. We use experimentally measured values of the growth rate for both forms of EN, with *r*_*pl*_ = 0.24 hr^−1^ for planktonic cells (i.e. clusters of size *n* = 1) and *r*_*agg*_ = 0.66 hr^−1^ for clusters (*n* ≥ 2). Growth occurs at every time step with *dt* being drawn from an exponential distribution with rate governed by the total overall reaction rate.

#### Aggregation

Aggregation involves a reaction of two clusters of sizes *n* and *m* combining to form a cluster of size *n* + *m*. Aggregation is taken to be size dependent with the rate of clusters *n, m* coming together given by:

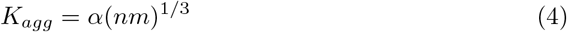

where 1/3 indicates that the collision of clusters is dependent on the cluster diameter and *α* = 10^−2.5^ hr^−1^. The clusters are randomly grouped in pairs at every time step and the rate of each aggregation reaction is computed.

#### Fragmentation

Fragmentation or breakup of a single cell from a cluster of size *n* occurs at a size dependent rate-

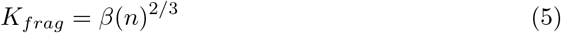

where 2/3 represents a surface area dependence, i.e. cells only on the surface of clusters can fragment. In our simulations, *β* is varied from 0.02 hr^−1^ to 1 hr^−1^ in steps of 0.025 hr^−1^. A value of 0.1 hr^−1^ corresponds to 10 cells fragmenting every hour from a cluster of size 1000.

#### Expulsion

The expulsion process involves removal of clusters from the intestine. Prior experimental work established that expulsion occurs on average every 10-12 hours, so the expulsion rate set to λ_*agg*_ = 0.1 hr^−1^ [72]. The expulsion rate of non-motile individuals occurs at a rate λ_*pl*_. The expulsion ratio 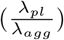 is varied logarithmically with 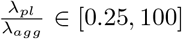.

#### Mono-association simulation

We first generate the initial mono-association cluster configuration for EN. Here, we keep *β* fixed at *β* = 0.01 hr^−1^, since in mono-association we see rare fragmentation of EN and log_10_(*α*) = −2.5. The expulsion rates are λ_*pl*_ = λ_*agg*_ = 0.1 hr^−1^. Growth of individuals and clusters occurs at rates *r*_*pl*_ = 0.24 hr^−1^ and *r*_*agg*_ = 0.66 hr^−1^.

At a given time step, the rates of aggregation, fragmentation and expulsion for all possible cluster reactions are computed. The total reaction rate i.e. the sum of all rates determines the jump interval (*dt*) when the next reaction occurs. At a given time step, only one cluster reaction is randomly chosen, weighted by the rate. The chosen reaction is executed, following which (λ_*pl*_*dt*) individual cells are expelled. (Expulsion of individuals is treated deterministically, for computational speed; note that there are large numbers of individuals, so stochasticity is unlikely to be important.) Before proceeding to the next time step, all clusters undergo growth as described previously.

The simulation is carried out until *T* = 24 hrs, the time at which the EN population is invaded by the mutant AE-MB4 species. The configurations with large (*n >* 1000) clusters are saved as the initial configuration for the invasion simulation.

#### Invasion simulation

We simulate for different parameters of *β* ∈ [0.02, 1] hr^−1^ in steps of df=0.02 hr^−1^ and 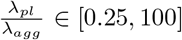. Within a simulation run, we choose 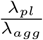 and start with the mono-association cluster size distribution. We evolve this distribution until T = 24 hrs (usually when the invasion abundance data is collected). All the processes i.e. growth, fragmentation, expulsion and aggregation are simulated exactly as in generating the initial cluster configuration. The total abundance along with final cluster sizes after 24 hrs are saved. The mean abundance from 500 fish for the full range of parameters of fragmentation rate and expulsion ratio are plotted in Figure 3 (D) with mean and variance of the data for each set of parameters provided in File S8.

The code for the simulation is provided on Github. (https://github.com/dsundarraman/zebrafish_*b*_*acterial*_*c*_*luster*_*d*_*ynamics.git*)

## Image analysis

### Identification of single cells and clusters in 3D images

Identification of single bacterial cells was done using a combination of machine learning and semi-automated approaches. A convolutional neural network previously trained on 3D images of single bacteria is used to classify potential objects as bacteria or noise blobs [16]. The same network architecture was used and the algorithm was trained on labeled datasets of AE-MB4 and EN cells. Identification of potential bacteria objects in voxels of size (30×30×10) pixels was done by using a local maxima finding algorithm in 2D and stitched in 3D. These potential objects were classified by the network and the classified output was curated manually.

For aggregate segmentation, the zebrafish gut was first segmented in 2D using the U-net algorithm which was previously trained on a hand-labeled dataset of zebrafish gut images [52]. After segmentation of the gut, bacterial clusters within the gut were segmented by applying a background threshold for each image of the z-stack for segmentation of bacterial aggregates. The total number of cells in an aggregate is determined by dividing the total fluorescence intensity from the segmented object by the median intensity of individuals in the image.

Once the net population in single cells and aggregates was ascertained, these were used to calculate the planktonic fraction, a ratio of the number of individuals to total abundance. To measure the cumulative cluster size distributions, p(n) and cluster size histograms depicting probabilities of finding a cell in an n-celled cluster, we performed a jackknife resampling on the dataset for each experiment to ascertain the mean and uncertainty in our measurements. The raw data files used for these is provided in File S2 and the calculated mean and uncertainties of these distributions is provided in File S4. The distributions shown in Fig 1 F and File S3 reflect pooled distributions with all clusters from all fish in an experiment.

### Normalized intensity profiles

After segmentation of the gut using U-net, we defined markers for the foremost point in the anterior in a 3D image. After applying a single background threshold for each 3D image, we computed the total intensity in this thresholded region along the intestinal axis. The normalized intensity profile over several fish is averaged and shown in Fig 1D, 5D and 6B. The raw data is provided in File S7.

### Growth rate measurement

To measure growth rates, we select consecutive 3D scans from fish, wherein aggregated or planktonic populations of bacteria are undisturbed and there is no influx or efflux of bacteria from fragmentation or expulsion respectively. We identify single cells and clumps using the image analysis pipeline described previously. The data is provided in File S5. An exponential fit to the abundance of single cells or clusters over time was used to determine the growth rate of cells and clusters respectively. The mean and the standard deviation from several fish are reported in Results.

## Author Contributions

All authors designed the research. DS, JVZK, and TJS carried out all experiments. DS analyzed the data with input from RP. DS and RP wrote the article with input from KG and TJS.

## Acknowledgments

We thank Rose Sockol, Ellie Melancon and the University of Oregon Zebrafish Facility staff for fish husbandry and Laura Taggart-Murphy for germ-free derivations of zebrafish larvae. This work was supported by the National Science Foundation under Award 1427957 and the National Institutes of Health under Awards P50GM09891 and P01GM125576. The University of Oregon Zebrafish Facility is supported by a grant from the National Institute of Child Health and Human Development (P01HD22486). The funders had no role in study design, data collection and analysis, decision to publish, or preparation of the manuscript.

## Supplemental Material

### 0.1 Supplemental Files

**File S1** Raw abundance data as CFUs from dissections and plating for all experiments. Each tab states the name of the experiment and each row is data from a single fish.

**File S2** Raw cluster size data for each fish from image analysis for all experiments. Each tab states the name of the experiment while rows indicate data from a single fish.

**File S3** Supplemental figures and text including information on model parameters and analysis.

**File S4** Mean frequencies and uncertainties from binned distributions showing the probability of being in an n-celled cluster for all experiments.

**File S5** Measured abundance of aggregated and planktonic populations of EN in the time lapse imaging experiments of fish in the AE-MB4 invasion of EN.

**File S6** Planktonic fraction over time from image analysis of three fish in the time series experiments of the AE-MB4 invasion of EN

**File S7** Raw data for measured intensity profiles from different experiments.

**File S8** Mean and uncertainties of the abundance from the cluster simulation for different paramaters of fragmentation rate and expulsion ratio 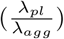. Each sheet name corresponds to the expulsion ratio parameter.

### 0.2 Supplemental Videos

Video titles are hyperlinks to videos uploaded on Vimeo.

**VideoS1** : Exemplar scans of 3D images showing the spatial distribution of AE (orange), EN (green) and AE-MB4 (magenta) in mono-association. Both AE and EN form large clusters in the midgut region of the larval zebrafish gut whereas planktonic cells of AE-MB4 are seen in the midgut and foregut region. Scale bar: 50 microns.

**VideoS2**: Exemplar scan of 3D image showing the spatial distribution of AE (orange) and EN (green) in the di-association experiment in the midgut region of the larval zebrafish gut. Co-aggregated populations of both species are prominent. Scale bar: 50 microns.

**VideoS3**: Exemplar scan of 3D image showing the spatial distribution of AE-MB4 (magenta) and EN (green) in the di-association experiment in the anterior and midgut region of the larval zebrafish gut. Single cells and small clusters of EN are surrounded by AE-MB4 cells. Scale bar: 50 microns.

**VideoS4**: Exemplar scan of 3D image showing the spatial distribution of AE-MB4 (magenta) and EN (green) in the anterior and midgut region of the larval zebrafish gut in the invasion experiment where EN colonized fish (5 dpf) are invaded by AE-MB4 (6 dpf). Single cells and a number of fragmented clusters of EN are surrounded by AE-MB4 cells and small clusters. Scale bar: 50 microns.

**VideoS5**: A series of maximum intensity projections of 3D images taken from time lapse imaging of a larval zebrafish gut initially colonized by EN 5 dpf and invaded by AE-MB4 6dpf, with imaging starting approximately 6 hrs post-invasion. The large EN aggregate initially fragments with individual cells being expelled from the gut as the AE-MB4 population grows and takes over. Scale bar: 100 microns.

**VideoS6**: A series of scans taken from time lapse imaging of a larval zebrafish gut initially colonized by EN (green) 5 dpf and invaded by AE-MB4 (not shown) 6dpf, with imaging starting approximately 6 hrs post-invasion. The scans show an initially aggregated population of EN (at 6 hr) breaking up over the course of time at 9.3 hr and 10 hr resulting in single cells in the midgut region. Scale bar: 50 microns.

**VideoS7**: Time lapse of a single plane showing single cells of non-motile EN (green) surrounded by AE-MB4 (magenta) in the AE-MB4 invasion of EN. An overall peristaltic flow slowly moves cells rightward. Scale bar: 20 microns.

**VideoS8**: Exemplar scan of 3D image showing the spatial distribution of AE (orange) and AE-MB4 (magenta) in the AE-MB4 invasion of AE in the midgut region of the larval zebrafish gut. Small clusters of AE are surrounded by AE-MB4 cells and clusters. Scale bar: 50 microns.

**VideoS9**: Exemplar scan of 3D image showing the spatial distribution of AE (orange and cyan) in the self invasion experiment in the midgut region of the larval zebrafish gut. Orange is the initial AE population which was invaded by isogenic AE in cyan. Co-aggregated populations of both orange and cyan are found and the orange cells are more dominant in the aggregate. Scale bar: 50 microns.

**VideoS10**: A series of maximum intensity projections of 3D images taken from time lapse imaging of a larval zebrafish gut initially colonized by AE 5 dpf and invaded by the AE-MB4 isolate 6dpf, with imaging starting approximately 6 hrs post-invasion.

Large aggregates of AE (orange) are expelled while smaller clumps of AE reside along a fast invading AE-MB4 (magenta) population. Scale bar: 100 microns.

**VideoS11**: Exemplar scan of a 3D image showing the a large aggregated cluster of EN (green) in the four species co-inoculation experiment in the midgut region of the larval zebrafish gut where the remaining three species (AC, PL and PS) are unlabelled. Scale bar: 50 microns.

**VideoS12**: Exemplar scan of a 3D image showing EN (green) and AE (orange) in the five species co-inoculation experiment (with AE) in the midgut region of the larval zebrafish gut where the remaining three species (AC, PL and PS) are unlabelled.

Co-aggregated populations of both species are shown. Scale bar: 50 microns. **VideoS13**: Exemplar scan of a 3D image showing EN (green) and AE-MB4 (magenta) in the five species co-inoculation experiment (with AE-MB4) in the midgut region of the larval zebrafish gut where the remaining three species (AC, PL and PS) are unlabelled. AE-MB4 resembles its mono-association phenotype, mostly consisting of single cells and smaller clusters while EN forms fragmented aggregates in the midgut region. Scale bar: 50 microns.

**VideoS14**: Exemplar scan of a 3D image showing EN (green) and AE-MB4 (magenta) in the five species invasion experiment in the anterior and midgut region of the larval zebrafish gut where the remaining four species (AC, AE, PL and PS) are unlabelled. The large aggregate of EN in the midgut persists while AE-MB4 co-aggregates in regions with EN. Scale bar: 50 microns.

**VideoS15**: Exemplar scan of a 3D image showing AE (orange) and AE-MB4 (magenta) in the five species invasion experiment in the midgut region of the larval zebrafish gut where the remaining four species (AC, EN, PL and PS) are unlabelled. The large aggregate of AE sustains itself in the presence of AE-MB4 and other species. Scale bar: 50 microns.

**VideoS16**: A series of maximum intensity projections of 3D images taken from time lapse imaging of a larval zebrafish gut initially colonized by EN (green) and the other 4 species (AC, AE, PL and PS-not shown) at 5 dpf and invaded by AE-MB4 at 6 dpf, with imaging starting approximately 6 hrs post-invasion. The AE-MB4 population is localized to the midgut region and the aggregate of EN shows slow fragmentation. Scale bar: 100 microns.

