## Supplemental S3: Text for "Disaggregation as an interaction mechanism among intestinal bacteria"

### Supplemental Materials

July 22, 2022

#### 1 Figures

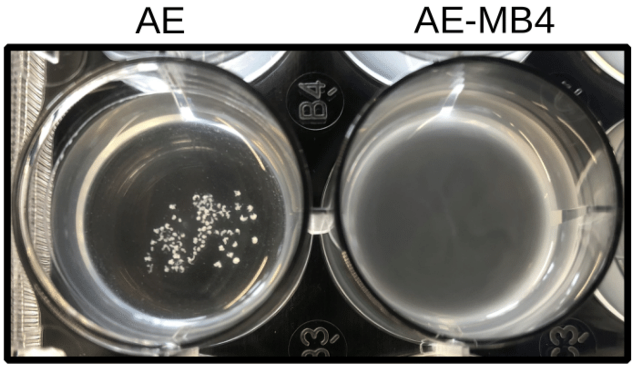


Figure 1: In vitro biofilm assay for AE (top panel) and AE-MB4 (bottom panel) in 0.4% GlcNAc solution. AE forms macroscopic aggregates in 0.4% GlcNAc (top-right) while AE-MB4 (bottom-right) is unable to form biofilms in the same medium.

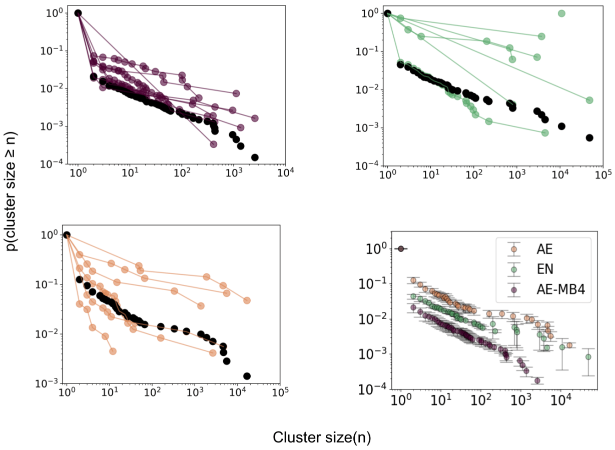


Figure 2: The cumulative cluster size distributions (p(cluster size $\geq$ n)) for (i) AE-MB4 (ii) EN (iii) AE and (iv) all three species in mono-association. The magenta, green and orange curves in (i), (ii) and (iii) are distributions calculated from clusters found in single fish in each mono-association experiment. The black curve in (i)-(iii) shows the pooled distribution calculated from clusters found in all fish. (iv) shows the pooled distribution for each of the species with all three showing power law behavior with slope m = 0.8 $\pm$ 0.1 for AE-MB4 and AE and m = 0.9 $\pm$ 0.2 for EN in the regime $n < 10^2$. Data are from N = 6, 7 and 8 fish for AE, EN and AE-MB4, respectively.

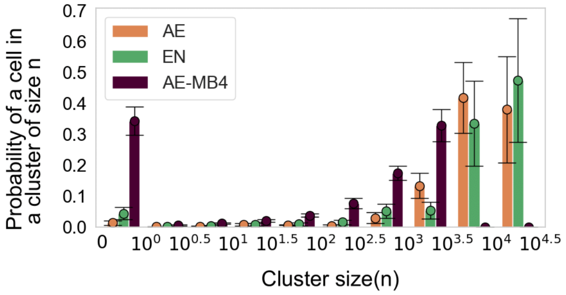


Figure 3: The probability of finding a cell in an n-cell cluster for AE (orange), EN (green) and AE-MB4 (magenta) in mono-association. Circles and error bars indicate the mean and uncertainties, calculated using jack-knife resampling. Tick marks indicate bin intervals, e.g. the three bars between 0 and $10^0$ correspond to EN, AE and AE-MB4 probabilities to be in clusters of size n = (0-1]. Data are from N = 6, 7 and 8 fish for AE, EN and AE-MB4, respectively.

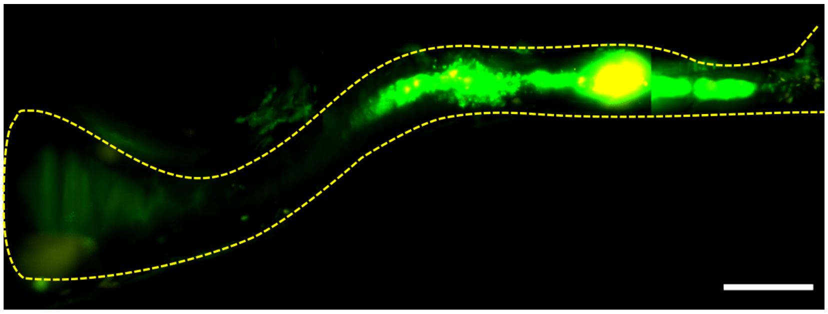

Figure 4: Maximum intensity projection of 3D image of a larval zebrafish gut colonized with EN (green) at 5dpf and challenged with AE (orange) at 6dpf. Overlapping populations are in yellow. Bar:100$\mu$m

### 2 Supplemental Text

### 3 Analysis summary

Raw data for all cluster sizes is provided in File S2. After identifying all clusters in the gut as described in methods main text, we use the list of clusters to generate (i) cumulative cluster size distributions and (ii) probability of finding a cell in an n-celled cluster.

#### 3.1 Cluster size distributions

For (i), we calculate the probability that a cluster will contain n cells for each fish. To calculate the mean and uncertainty of the distribution, we use jack-knife sampling, pooling clusters from all fish, except one and calculating the cumulative cluster size distribution. These distributions are shown in Figure 1 and Supplemental Figure S2.

The power law fits were done using maximum likelihood estimation using the python version of the code in https://aaronclauset.github.io/powerlaws/ provided in https://github.com/jeffalstott/powerlaw. For the fit, only the initial part of the distribution, n¡100 was used. We used jack-knife sampling on the dataset to compute the mean and variance of the power law exponent.

#### 3.2 Probability of being in an n-celled cluster

After obtaining cluster sizes from different fish (File S2), we perform a similar subsampling of the data, leaving out a single fish and binning clusters from all the remaining fish in fixed logarithmic sized bins and compute the probability of finding a cell in each bin by dividing the total number of cells in the bin to the net abundance of all clusters. We then calculate the mean and uncertainty of the frequencies of the various subsamples for each bin. These are provided in File S3.
