## Supplementary figures and images for "Disaggregation as an interaction mechanism among intestinal bacteria"

### Figure S1

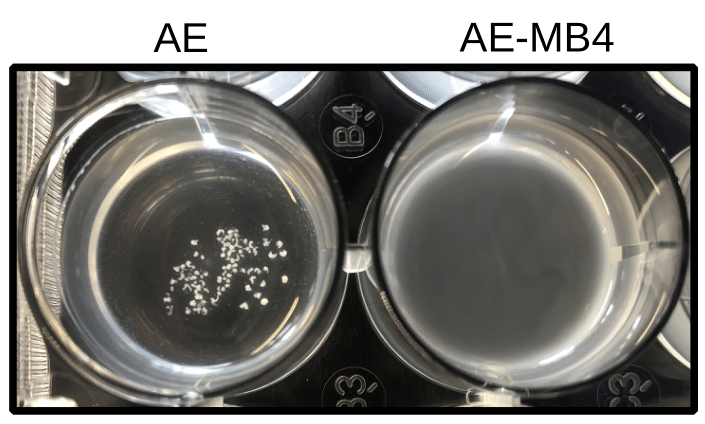

### Figure S2

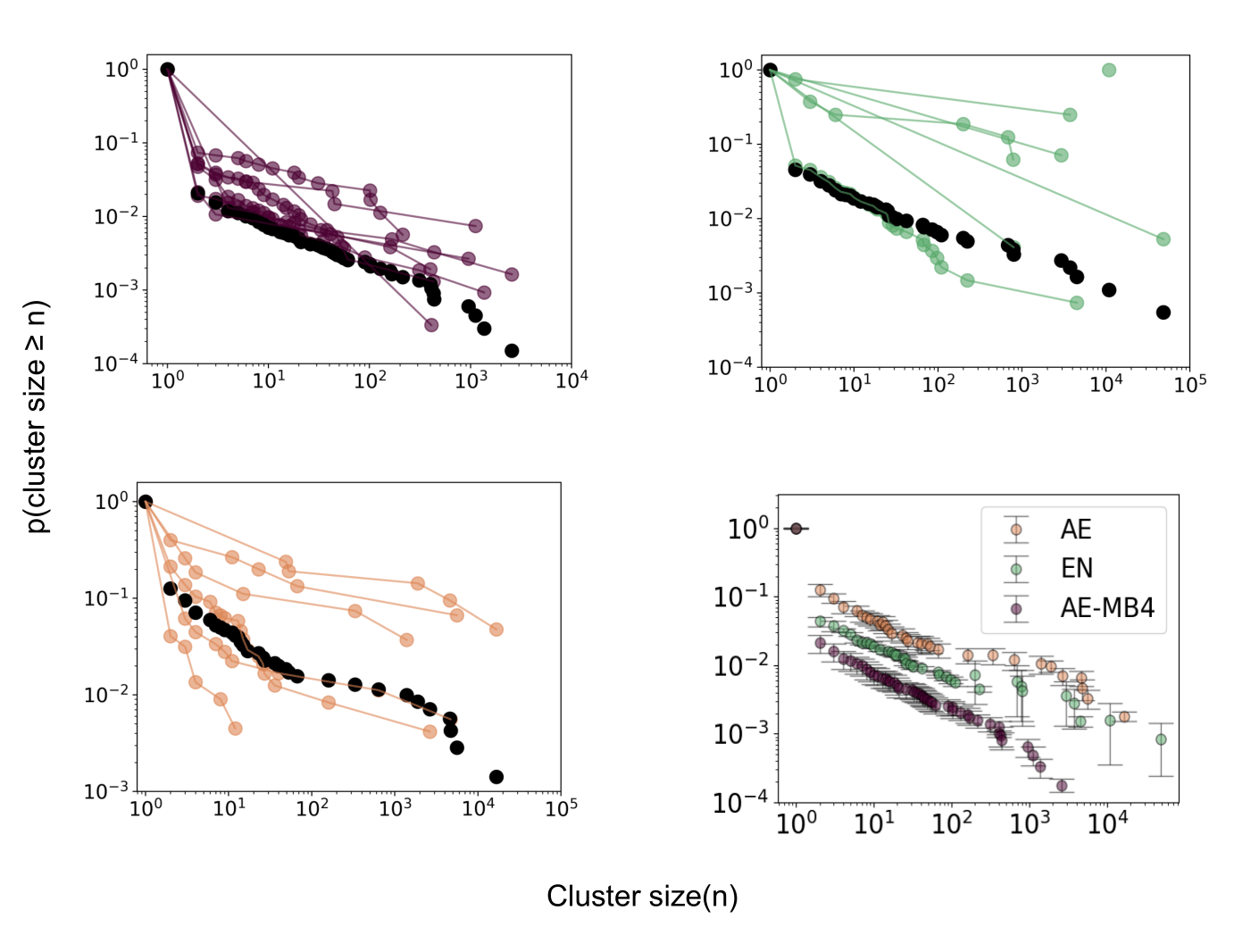

### Figure S3

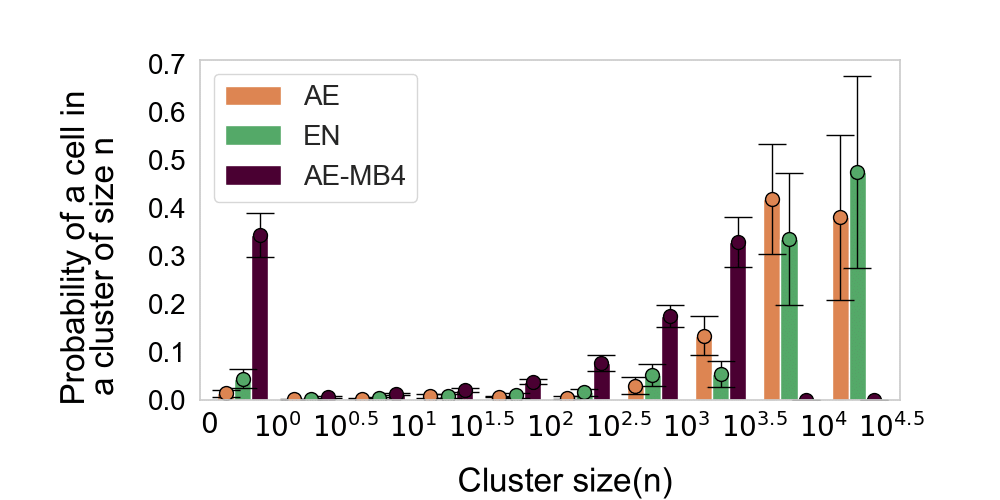

### Figure S4

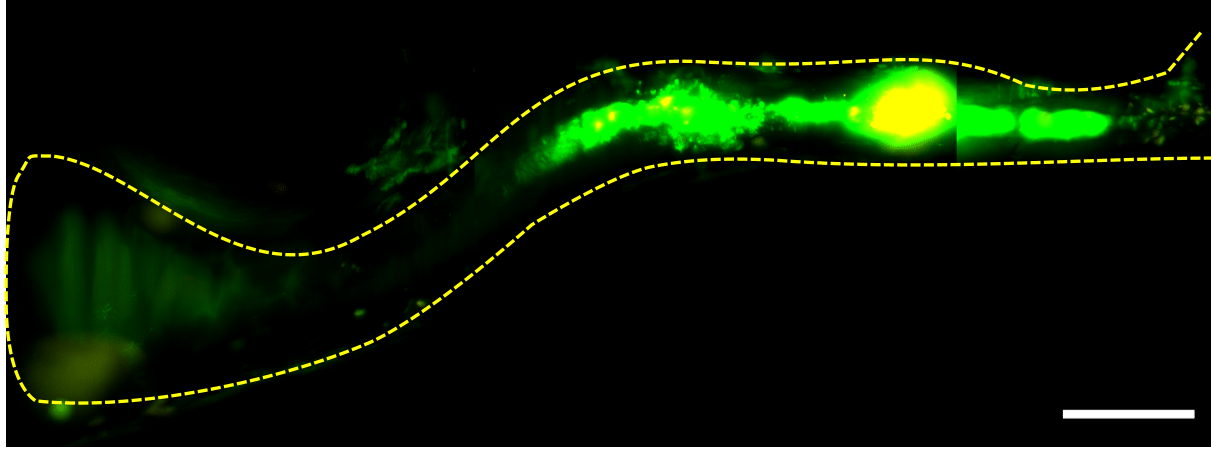
